# Machine Learning-Driven Drug Repurposing for KRAS G12C and KRAS G12D Inhibition

**DOI:** 10.1101/2025.05.16.654410

**Authors:** Gianluca Fuschi, Julia St. Germain, David Bebensee, Christophe Moawad, Arina Aladysheva, Ashraf Mohamed, Eiman Elwakeel, Bernard R. Brooks, Muhamed Amin

## Abstract

KRAS is a predominant oncogenic driver across multiple cancers, long considered “undruggable” due to its high nucleotide affinity and lack of classical binding pockets. Although recent advances have led to covalent inhibitors like Sotorasib and Adagrasib for the KRAS G12C mutation, effective therapies for other common variants—most notably KRAS G12D, which is highly prevalent in aggressive pancreatic cancers—remain limited. In this study, we employ machine learning to identify potential inhibitors for both KRAS G12D and G12C by screening FDA-approved compounds from the ChEMBL database. Random Forest and Neural Network models were trained on binding affinity data from three BindingDB datasets: wild-type KRAS GTPase, KRAS G12C, and KRAS G12D. Our models identified high-affinity candidates including anti-cancer kinase inhibitors (e.g., Cobimetinib, Gilteritinib) as well as drugs from other categories (e.g., Bromocriptine, Cefepime). By incorporating atomic hybridization as a feature, we aim to capture the effects of induced polarization, potentially improving prediction accuracy. Despite modest agreement across models, our approach highlights several promising candidates—such as Acalabrutinib, which has prior evidence of G12C activity—for further investigation. These results offer a data-driven foundation for experimental validation and the future development of targeted KRAS therapies.

## 1 INTRODUCTION

Despite decades of effort, targeting RAS-driven cancers remains an unmet need: activating RAS mutations occur in roughly 20% of all tumors [1], and KRAS alone drives 76% of these cases. KRAS is a small GTPase that cycles between an active GTP-bound form and an inactive GDP-bound form under the control of GEFs and GAPs [2]. Mutations impair its intrinsic GTP hydrolysis, locking KRAS in the signaling-competent state and driving RAF–MEK–ERK activation, tumor progression, and therapy resistance [3].

KRAS comprises a conserved G-domain, which binds GTP/GDP, and a C-terminal hypervariable region (HVR) that anchors it to the membrane via farnesylation at the CAAX motif [2, 4]. Within the G-domain, the P-loop (Walker A) and the switch-I (aa 30–38) and switch-II (aa 60–76) regions form the interface for GAPs, GEFs, and downstream effectors [5, 6].

Nearly 98% of KRAS mutations arise in codons 12, 13, or 61—altering the P-loop or switch-II and locking KRAS in its GTP-bound form [7, 8]. Until recently considered “undruggable,” the discovery of a switch-II pocket enabled the development of covalent G12C inhibitors [9]. Sotorasib and adagrasib, which irreversibly bind the SII-P pocket of KRAS G12C and trap it in the GDP-state, received accelerated FDA approval in 2021 and 2024, respectively, for NSCLC [10, 11].

These inhibitors are, however, limited to KRAS G12C. To overcome this limitation, pan-KRAS and even pan-RAS compounds, like BI-2852 and RMC-7977, respectively, have been developed and evaluated for their binding capabilities to other KRAS mutations, highlighting the potential for broader therapeutic applications[12, 13].

By contrast, KRAS G12D—prevalent in pancreatic ductal adenocarcinoma, the second leading cause of cancer death—is refractory to G12C strategies [14]. The aspartate substitution disrupts GAP-mediated hydrolysis by reorienting switch-I/II and altering the active site charge [5]. Several G12D-specific inhibitors are now in early clinical development, including the non-covalent MRTX1133 and the cyclophilin-A–assisted RMC-9805, as well as ASP3082 and HRS-4642; others remain preclinical (TH-Z835, JAB-22000, ERAS-4693/-5024) [5].

Machine learning (ML) has transformed drug discovery—accelerating target validation, biomarker identification, and digital pathology analyses in clinical trials [15]. Drug repurposing, in particular, leverages known safety profiles to compress timelines compared to de novo design [16, 17].

Prior repurposing efforts against KRAS include ML-driven docking and MD studies that nominated EGFR inhibitors (afatinib, neratinib, zanubrutinib) for G12C [18], as well as screens identifying antibiotics and anti-inflammatories (capreomycin, cefdinir, cortisone) for G12C/G12S [19]. For G12D, virtual screening and XP docking studies have proposed hydroxyzine, zuclopenthixol, fluphenazine, and doxapram as candidates [20].

Here, we integrate ML models—enhanced with atomic-hybridization and induced-polarization features—with molecular docking and MD simulations to screen all FDA-approved drugs against KRAS G12D. We then benchmark our top hits against literature-reported compounds to validate and extend previous findings.

## 2 METHODS

The set of potential inhibitors targeting carcinogenic KRAS mutations was extracted from three datasets using two ML approaches. These datasets include wild-type KRAS and two mutant forms—G12C and G12D. By analyzing these datasets separately, we aimed to identify overlapping predicted molecules, under the assumption that compounds common to all three are less likely to exhibit specificity toward the mutant variants and thus may be less effective inhibitors. The use of two different ML models on each dataset allowed us to assess the internal consistency of predictions; compounds identified by both models within a single dataset are considered stronger candidates. In total, six lists of predicted molecules were generated and screened for promising candidates based on model agreement and dataset specificity. The top candidates are highlighted in the discussion for further experimental investigation.

### 2.1 Data preparation

The training data for the machine learning (ML) model were obtained from the ChEMBL [21] and BindingDB [22] databases, both of which offer extensive coverage of bioactivity data. These datasets were filtered based on the target protein: GTPase KRAS (from ChEMBL), and the KRAS variants with G12C and G12D mutations (from BindingDB). A previously developed computational drug discovery framework was employed to train ML models using the ChEMBL bioactivity data [23]. Only molecules with experimentally recorded binding affinities (IC50 values) to their respective KRAS targets were retained. Molecular descriptors were extracted using the RDKit Python library [24], and additional features were generated based on the count of atoms in each possible hybridization state, following an adaptation of a method introduced in a recent study on induced polarization in drug discovery [25]. The target variable for the ML models was the IC50 binding affinity of each molecule to either GTPase KRAS, KRAS G12C, or KRAS G12D, measured in nanomolar units. The IC50 values were standardized using a logarithmic transformation to convert them into pIC50 values. To prevent bias due to differing numerical scales, both the input features and the target variable were normalized using the MinMaxScaler from Scikit-learn [26].

For the three datasets considered in this study, only the FDA-approved compounds classified as “Small molecules”, “Approved” for max stage, and “0; 1” for Rule of Five (RO5) violations are considered. The molecules were initially screened using Lipinski’s RO5 to eliminate compounds lacking drug-like properties and to increase the likelihood of bioactivity [27]. Moreover, molecules with the same SMILES code and IC50 value were removed. Due to the reduced sizes of the datasets, a large number of features can lead to overfitting—where the model learns noise instead of underlying patterns—and also increase training time [28, 29]. Therefore, variance threshold and univariate feature selection methods were applied to retain the most influential features using the Scikit-learn library [26].

### 2.2 Machine Learning models

The Machine Learning libraries used for this study are Scikit-learn [26] for the Random Forest (RF) and PyTorch [30] for the Neural Network (NN). To assess the model capability to generalize to unseen data, the best RF candidate model is optimized using k-Fold cross-validation with k is set to 15 folds. The parameters of the RF model were set to a maximum tree depth of 20 and a minimum leave size of 10. On the other hand, the NN is inherited from a recently developed model [25]. A model is optimized for IC50 prediction exploiting the atomic hybridization features. Similarly, the NN model was trained using K-Fold cross-validation with K is set to 5 folds. The NN is built with 4 hidden layers 256 nodes each, and trained for 200 epochs. The full model architecture is detailed in [25] The accuracy of the models was assessed using the R^2^ score and Mean Squared Error.

## 3 RESULTS

We trained Random Forest (RF) and Neural Network (NN) models on bioactivity data for wild-type KRAS and its G12C and G12D mutants to identify FDA-approved drugs with high predicted binding affinity.

1. For GTPase KRAS dataset: R^2^ score is 0.663 (MSE: 0.0203) for NN model and 0.642 (MSE: 0.2063) for RF model.
2. For G12D dataset: R^2^ score is 0.676 (MSE: 0.0064) for NN model and 0.642 (MSE: 0.2063) for RF model.
3. For G12C dataset: the R^2^ score is 0.934 (MSE: 0.0129) for NN model and 0.939 (MSE: 0.2445) for RF model.

Figure 1 shows that both models achieve similar R^2^ values, but the NN exhibits greater overfitting, likely because its number of trainable parameters is large relative to the size of the datasets.

**Figure 1.**
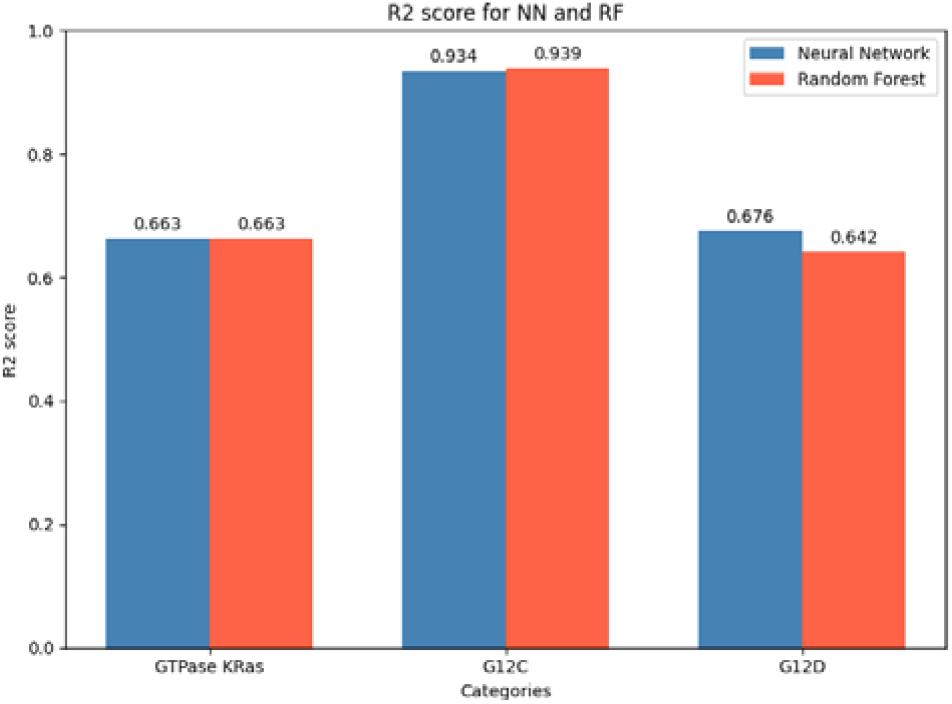
R^2^ score comparison between different Random Forest and Neural Network models for each dataset.

The top 10 predicted molecules for each dataset and model are add to the SI materials (Tables S1 to S6). Each table includes the ChEMBL ID, molecule name, drug class, mechanism of action, and predicted binding affinity. Figure 2 illustrates the variation in model performance across the three datasets—GTPase KRAS, G12C, and G12D—which is partly due to differences in dataset sizes. In particular, the G12D dataset has more entries, enabling more effective training and evaluation of both the NN and RF models. To better understand the influence of individual molecular features, permutation importance was used to identify the top 10 most impactful features, as shown in Figure 3. These features represent the key molecular descriptors driving the models’ binding affinity predictions to KRAS targets.

**Figure 2.**
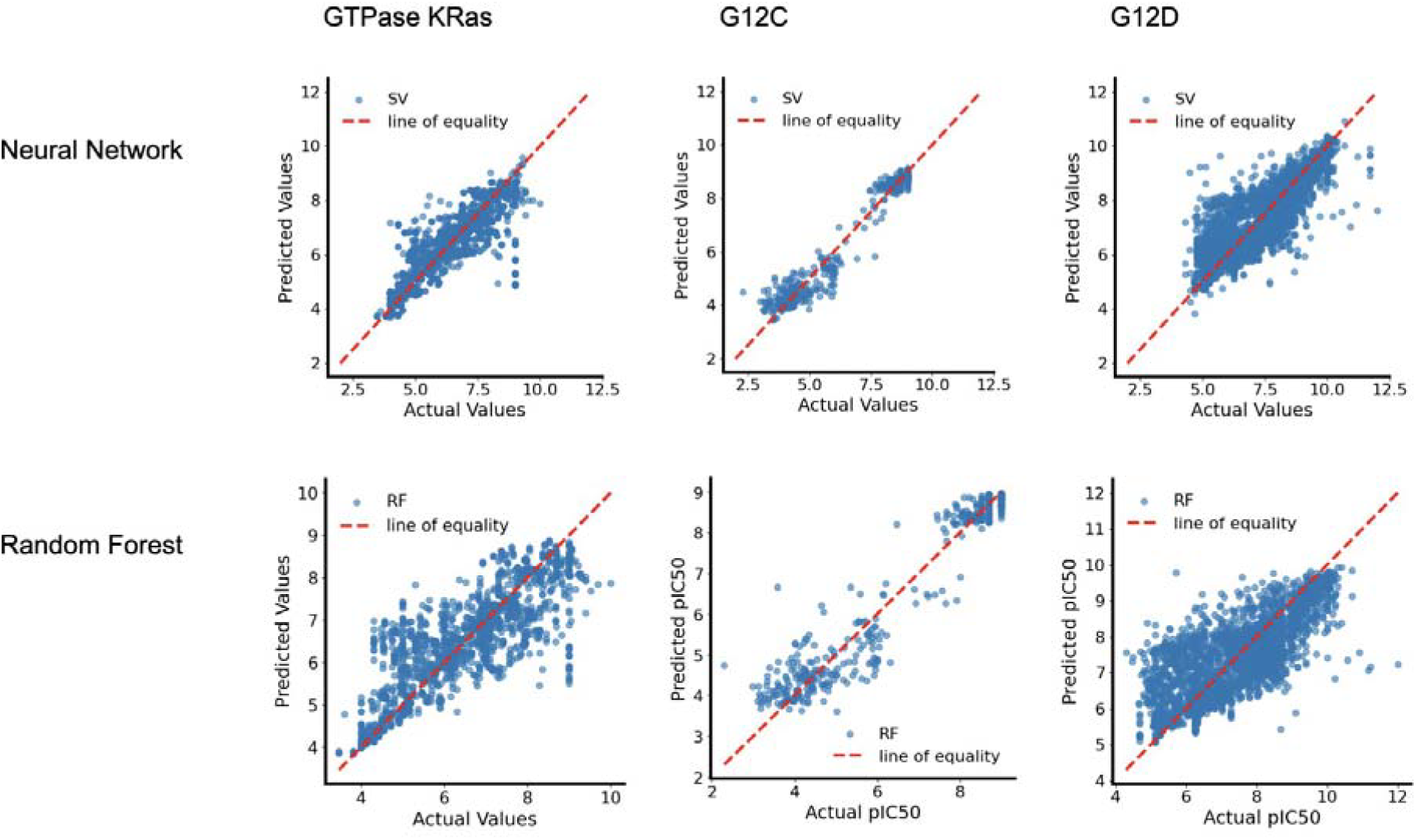
Actual vs Predicted values for Random Forest and Neural Network models for each dataset.

**Figure 3.**
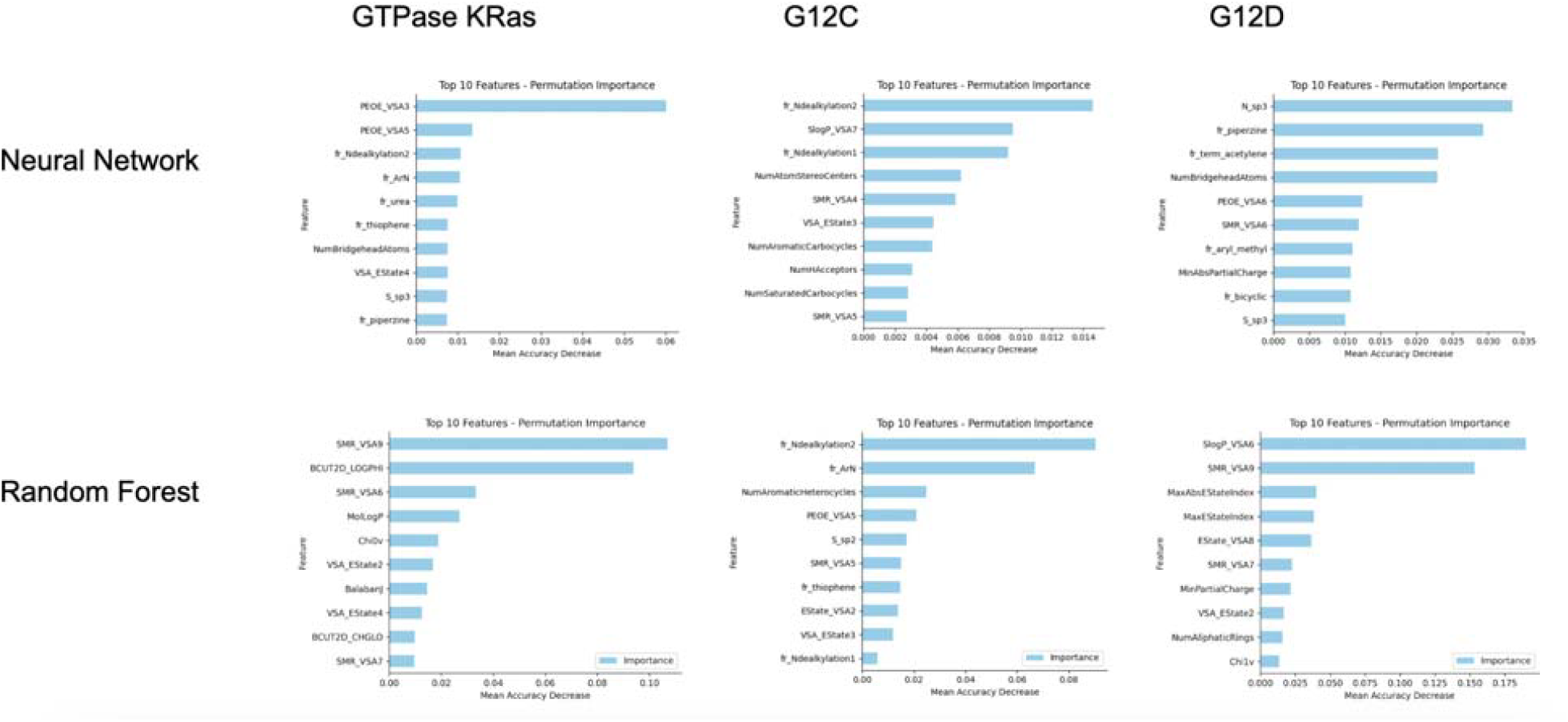
Feature importance for each dataset for NN and RF models.

## 4 DISCUSSION

This study highlights the molecular features that most significantly influence the prediction of binding affinity to KRAS. Among the most predictive features across all models are S_*sp*3_ and N_*sp*3_, which represent the counts of sulfur and nitrogen atoms in *sp*^3^ hybridization states. In addition, hybrid electronic-topological descriptors—such as PEOE_*VSA*_ (partial charge-weighted surface area), SMR_*VSA*_ (molar refractivity-weighted surface area), and EState_*VSA*_ (electrotopological state-weighted surface area)—emerge as key predictors. These features are closely linked to atomic hybridization and electronic structure, emphasizing the critical role of induced polarization and electronic configuration in KRAS-ligand molecular recognition.

Although all Neural Network (NN) models demonstrated relatively high predictive accuracy, their top-10 predicted molecules showed no overlap with those of the Random Forest (RF) models on the same datasets, suggesting limited agreement between the two approaches. This discrepancy may be partially attributed to differences in feature importance, as shown in Figure 3. However, when the comparison is extended to the top-30 predicted molecules, some overlap is observed—specifically, two shared compounds for the G12C dataset and three for the GTPase KRAS dataset—while the G12D dataset still shows no overlap (Tables S7 and S8). This divergence in predictions suggests that the models are not relying on straightforward correlations between individual features and the target variable. Instead, they are likely capturing more complex, nonlinear patterns in the data, with important features emerging through intricate interactions rather than direct associations.

In contrast, some predicted compounds (Tables S1–S6) appeared across multiple KRAS variant datasets (e.g., G12C, G12D, and GTPase), suggesting potential binding to conserved regions of the KRAS protein rather than mutation-specific sites. For example, Gilteritinib Fumarate and Telotristat Etiprate were predicted by the NN model for both G12C and GTPase datasets, while Rescinnamine appeared in G12D RF and GTPase NN predictions. Although these compounds demonstrated high predicted pIC50 values, their presence in both wild-type and mutant predictions may indicate a lack of selectivity, which reduces their potential as mutation-specific inhibitors. Rimegepant Sulfate, a CGRP receptor antagonist used for migraine treatment, was predicted in both G12C RF and G12D NN datasets. This could imply either a lack of mutation specificity or possible efficacy as a pan-KRAS inhibitor. However, considering its known mechanism of action and relatively low predicted pIC50, its potential for KRAS targeting is likely limited.

Unexpectedly, the GTPase RF model predicted both Adagrasib and Sotorasib—two well-known KRAS G12C inhibitors—which was not expected, given that WT KRAS used in the dataset lacks the cysteine mutation necessary for those compounds to bind. That might indicate that features that are not impacted by the mutated cysteine residue are important for determining the probability of the small molecules binding to KRAS. Conversely, none of the models that were trained on KRAS G12C dataset predicted any of these two drugs, making their results less reliable.

Overall, computational analysis identified several drug molecules predicted to interact with GTPase and KRAS mutants (G12C/G12D). Notably, multiple anti-cancer agents were prominent in addition to Sotorasib and Adagrasib, including Cobimetinib (8.95), Gilteritinib Fumarate (7.75), Ivosidenib (7.37), Clofarabine (7.27), Apalutamide (7.21), Trimetrexate Glucuronate (7.19), Acalabrutinib Maleate (6.96), Cladribine (6.9), Cobimetinib Fumarate (6.66) and Copanlisib (5.45). Interestingly, the highest pIC50 scores were shared between kinase inhibitors (Cobimetinib, Gilteritinib Fumarate), showing that these particular types of inhibitors are preferred by the algorithms. Notably, Gilteritinib Fumarate—a tyrosine kinase inhibitor used in acute myeloid leukemia treatment—had one of the highest pIC50 (8.89 in G12C NN, 7.75 in GTPase NN), yet its presence across both mutant and GTPase KRAS types implies that it is not mutant-specific.

Other drug classes identified through the models also warrant further investigation. For example, Bromocriptine, predicted as a top candidate by the KRAS G12D RF model (predicted pIC50: 7.86), has previously been reported to inhibit drug-resistant tumor cells in a separate drug-repurposing study [31]. Similarly, Cefepime, an antibiotic predicted by the KRAS G12C RF model (predicted pIC50: 7.56), has demonstrated tumor-depleting activity in preclinical studies [32]. In addition, several analgesic compounds were also highlighted, including codeine sulfate, morphine sulfate, and sufentanil. Prior research suggests that sufentanil may inhibit tumor growth [33], while morphine has been implicated in modulating cancer cell proliferation and survival pathways [34]. These findings suggest that even non-oncological drugs predicted by the models may have repurposing potential and merit further biological validation.

Moreover, KRAS G12C RF identified Acalabrutinib - an anti-cancer drug used for lymphoma treatment, which was also mentioned as a potential active inhibitor for KRAS G12C in a previous in-silico drug-repurposing study, requires further attention [18]. On the other hand, covalent docking of Acalabrutinib in that study showed that it bond with the cysteine 12 residue in a place different than Sotorasib and Adagrasib, particularly, on the outside of the allosteric pocket, making it less likely to successfully act as an inhibitor.

Some of the predicted molecules are less likely to act as an inhibitor due to their well-known mechanisms of action - such as Iobenguane Sulfate, which is an imaging agent or Nalmefene Hydrochloride, which is used to manage opioid overdose. Moreover, Methapyrilene Fumarate was recently deemed acutely toxic and revoked, likely meaning that the ChEMBL dataset of the FDA-approved molecules was not up to date [35].

This study has several limitations, the main being inherent in its computational nature. The dataset used for this study might have certain biases - ranging from repetition of components to the differences in the experimental procedures used to obtain the data. For instance, as we discovered, the ChEMBL dataset did not mark Methapyrilene as highly-toxic, suggesting that it might be incomplete or outdated, might influence our prediction models. Moreover, while computational models provide valuable insights and can efficiently narrow down potential drug candidates, they may not fully capture the complexity of biological interactions. This highlights the importance of incorporating uncertainty estimates into predictive modeling workflows to better assess confidence in the predictions and prioritize molecules for experimental validation. By integrating uncertainty quantification, such models can become more robust, interpretable, and ultimately more reliable in guiding drug discovery efforts. While some of the effects could have been mitigated by adding molecular docking to the study, computational predictions still require experimental validation.

## 5 CONCLUSION

This study employed Machine Learning (ML) to identify FDA-approved compounds with potential high binding affinity to wild-type GTPase KRAS, as well as KRAS G12C and G12D. While the models demonstrated reasonable predictive accuracy, the lack of strong consensus between different approaches and datasets highlights the challenges in computational drug repurposing. The discrepancies with previously reported inhibitors may arise from differences in dataset composition, model selection, or docking methodologies. Despite these limitations, the study provides a foundation for further investigation, particularly for compounds like Acalabrutinib, which was also predicted to be an inhibitor by a different study, as well as Cobimetinib, Gilteritinib Fumarate, Bromocriptine and Cefepime. Future work should focus on experimental validation and refining computational models incoprating uncertainness estimates to improve KRAS inhibitor discovery. Integrating dynamic simulations and advanced ML techniques could enhance prediction reliability, ultimately aiding in the development of targeted therapies for KRAS-induced cancers.

## 6 AUTHOR INFORMATION

### 6.1 Authors

Gianluca Fuschi^†^, Julia St. Germain^†^, Arina Aladysheva^†^, David Bebensee^†^, Christophe Moawad^†^, Ashraf Mohamed, Eiman Elwakeel, Bernard R. Brooks, Muhamed Amin

### 6.2 Author Contributions

## 6.3 Acknowledgments

For their contribution to this work, we thank Aniela Hofmokl, Claudia Jara Soto, Liv Mueller, Jade Howard, Merel Wiesenekker, Fiona, Chun Lei Koenders, Maxime Heinsbroek, Quinten Post, Victor Giustini Perez, Roos De Jong, Hanna Maouche, Fiona Majumdar.

