## Supporting Information for "Machine Learning-Driven Drug Repurposing for KRAS G12C and KRAS G12D Inhibition"

**^1^Department of Sciences, University College Groningen, University of Groningen, 9718 BG Groningen, The Netherlands**

**^2^Department of Chemistry, University of Texas at Austin, Austin, Texas 78712, USA**

**^3^Max Planck Institute for Heart and Lung Research, Bad Nauheim, Germany**

**^4^Institute for Lung Health (ILH), Justus Liebig University, Giessen, Germany**

**^5^Laboratory of Computational Biology, National Heart, Lung and Blood Institute, National Institutes of Health, Bethesda, Maryland 20892, USA**

###

**Supporting Information**

Github repository that contains the data and code use for predicting the binding affinity.
URL: <https://github.com/juliastgermain/ML_for_KRAS_G12D_Inhibitors>’

**Table S1**

|  | Gtpase Random Forest |  |
| --- | --- | --- |
| CHEMBL ID | Molecule Name | Predicted PiC50 |
| CHEMBL4594350 | ADAGRASIB | 8.106427137 |
| CHEMBL2359966 | QUININE SULFATE | 7.497035617 |
| CHEMBL3707183 | QUINIDINE SULFATE | 7.497035617 |
| CHEMBL4297091 | CODEINE SULFATE | 7.425687233 |
| CHEMBL2361370 | PENBUTOLOL SULFATE | 7.419709268 |
| CHEMBL3989958 | IVOSIDENIB | 7.372708472 |
| CHEMBL3187985 | APOMORPHINE HYDROCHLORIDE | 7.257022141 |
| CHEMBL2103744 | MORPHINE SULFATE | 7.225236355 |
| CHEMBL3183409 | APALUTAMIDE | 7.217008093 |
| CHEMBL4535757 | SOTORASIB | 7.173295985 |

**Table S2**

G12C Random Forest

| CHEMBL ID | Molecule Name | Predicted PiC50 |
| --- | --- | --- |
| CHEMBL1200962 | CEFEPIME HYDROCHLORIDE | 7.5636268 |
| CHEMBL186 | CEFEPIME | 7.560583672 |
| CHEMBL3187246 | METHAPYRILENE FUMARATE | 7.259904943 |
| CHEMBL1201163 | SUFENTANIL CITRATE | 7.173398563 |
| CHEMBL2364629 | RIMEGEPANT SULFATE | 7.0659574 |
| CHEMBL658 | SUFENTANIL | 7.040832279 |
| CHEMBL2105458 | THENALIDINE | 7.039106467 |
| CHEMBL534 | KETOTIFEN | 7.023874216 |
| CHEMBL1633 | KETOTIFEN FUMARATE | 7.01436391 |
| CHEMBL4594293 | ACALABRUTINIB MALEATE | 6.963268857 |

**Table S3**

G12D Random Forest

| CHEMBL ID | Molecule Name | Predicted PiC50 |
| --- | --- | --- |
| CHEMBL1200503 | BROMOCRIPTINE MESYLATE | 7.863023 |
| CHEMBL493 | BROMOCRIPTINE | 7.804625 |
| CHEMBL1668 | RESCINNAMINE | 7.762681 |
| CHEMBL1200722 | PIPECURONIUM BROMIDE | 7.769046 |
| CHEMBL4297066 | ELIGLUSTAT TARTRATE | 7.669139 |
| CHEMBL772 | RESERPINE | 7.604386 |
| CHEMBL1200648 | ROCURONIUM BROMIDE | 7.574692 |
| CHEMBL1737 | SILDENAFIL CITRATE | 7.55759 |
| CHEMBL2105891 | PHYSOSTIGMINE SULFATE | 7.545276 |
| CHEMBL1201206 | PIPECURONIUM | 7.422963 |

**Table S4**

Gtpase Neural Network

| CHEMBL ID | Molecule Name | Predicted PiC50 |
| --- | --- | --- |
| CHEMBL2146883 | COBIMETINIB | 8.95349 |
| CHEMBL3301603 | GILTERITINIB FUMARATE | 7.759499 |
| CHEMBL1200678 | ATAZANAVIR SULFATE | 7.5652604 |
| CHEMBL1668 | RESCINNAMINE | 7.5329623 |
| CHEMBL3348963 | TELOTRISTAT ETIPRATE | 7.1994147 |
| CHEMBL1201244 | ROCURONIUM | 6.9268174 |
| CHEMBL6966 | VERAPAMIL | 6.3693757 |
| CHEMBL282724 | SITAXENTAN | 5.7964606 |
| CHEMBL2040682 | CICLESONIDE | 5.6234417 |
| CHEMBL3218576 | COPANLISIB | 5.4535313 |

**Table S5**

G12C Neural Network

| CHEMBL ID | Molecule Name | Predicted PiC50 |
| --- | --- | --- |
| CHEMBL3301603 | GILTERITINIB FUMARATE | 8.891777 |
| CHEMBL1088977 | ADEMETIONINE | 7.537722 |
| CHEMBL1750 | CLOFARABINE | 7.2668715 |
| CHEMBL2218878 | TRIMETREXATE GLUCURONATE | 7.1885242 |
| CHEMBL3348963 | TELOTRISTAT ETIPRATE | 7.1353893 |
| CHEMBL3989695 | REGADENOSON | 6.97351 |
| CHEMBL1619 | CLADRIBINE | 6.905672 |
| CHEMBL1167 | SPECTINOMYCIN | 6.6144476 |
| CHEMBL278623 | MACIMORELIN | 6.5715756 |
| CHEMBL1753 | CLINDAMYCIN | 6.3678656 |

**Table S6**

G12D Neural Network

| CHEMBL ID | Molecule Name | Predicted PiC50 |
| --- | --- | --- |
| CHEMBL3989933 | ETRASIMOD ARGININE | 6.7378364 |
| CHEMBL1201327 | ACETRIZOIC ACID | 6.7078786 |
| CHEMBL1624 | LEVOTHYROXINE | 7.682719 |
| CHEMBL559 | DEXTROTHYROXINE | 7.6716547 |
| CHEMBL2364607 | COBIMETINIB FUMARATE | 6.664458 |
| CHEMBL3989511 | IOBENGUANE SULFATE I 131 | 6.5087504 |
| CHEMBL3989523 | IOBENGUANE SULFATE I 123 | 6.5047216 |
| CHEMBL2364629 | RIMEGEPANT SULFATE | 6.412818 |
| CHEMBL5315055 | NALMEFENE HYDROCHLORIDE DIHYDRATE | 6.33625 |
| CHEMBL3544986 | PERINDOPRIL ARGININE | 6.3038526 |

Predicted molecules that overlap between NN and RF, after the top 10 prediction.

**Table S7**

G12C Overlapping molecules

| CHEMBL ID | Molecule Name | NN predicted PiC50 | RF predicted PiC50 |
| --- | --- | --- | --- |
| CHEMBL186 | CEFEPIME | 6.697 | 7.477 |
| CHEMBL1200962 | CEFEPIME HYDROCHLORIDE | 6.609 | 7.477 |

**Table S8**

GTPase KRAS Overlapping molecules

| CHEMBL ID | Molecule Name | NN Predicted PiC50 | RF Predicted PiC50 |
| --- | --- | --- | --- |
| CHEMBL3301603 | GILTERITINIB FUMARATE | 7.275 | 6.915 |
| CHEMBL3348963 | TELOTRISTAT ETIPRATE | 6.900 | 6.984 |
| CHEMBL1668 | RESCINNAMINE | 7.439 | 7.072 |
